# Estimation of metabolite levels in cheese from microbial gene expression

**DOI:** 10.64898/2026.04.04.716484

**Authors:** Anis Mansouri, Rina Mekuli, Dominique Swennen, Francesco Durazzi, Daniel Remondini

**Affiliations:** Department of Physics and Astronomy, University of Bologna, Italy; l’Institut National de Recherche pour l’Agriculture, France

## Abstract

Characterizing aroma and flavours generated during cheese production is of high relevance for the food industry. A deeper comprehension of flavour generation can be achieved by understanding the role of microbial population governing milk processing, and in particular their metabolic activity governed by gene expression.

In this work we considered two independent experiments in which gene expression of the microbial population involved in cheese processing is sampled, together with final volatile products quantification. We estimated the final volatile compound profile from the measured metatranscriptomic expression by using machine learning with two different strategies for model training and validation, and we were able to associate specific biochemical pathways to the identified gene signatures.

## Introduction

Cheese is a dairy product that has been produced since the earliest civilizations some 8000 years ago during the “Agricultural Revolution” (Patrick, 2000). Its production spread throughout Europe and the Middle East and later to North and South America and Oceania. Nowadays, there are at least 1000 cheese varieties produced all over the world (Sandine & Elliker, 1970). The conversion of fresh milk into cheese involves different microbial species including bacteria, yeasts and moulds which perform the three major pathways constituting the biochemistry of cheese fermentation and ripening: (1) metabolism of residual lactose and of lactate and citrate (primary reactions), (2) lipolysis and fatty acid metabolism (3) proteolysis and amino acid catabolism (secondary reactions). These biochemical processes result in the development of flavor and texture characteristics of the cheese. For instance, Lc. lactis ssp. lactis and Leuconostoc spp metabolize citrate to diacetyl in the presence of fermentable sugar during manufacture and early ripening. Diacetyl contributes to the flavor of Dutch-type cheeses and possibly Cheddar also. The CO2 produced is responsible for the small eyes characteristic of Dutch-type cheeses. The metabolism of fatty acids in cheese by Penicillium spp produces n-methyl ketones which dominate the taste and aroma of blue cheese (Patrick, 2000).

Cheese flavor and aroma are among the main properties that determine cheese’s quality and influence consumers’ preferences (Biolatto et al., 2007; Gómez et al., 2006; Korel & Balaban, 2002). For that reason, one of the main issues for the dairy industry is monitoring and characterizing cheese’s composition, aroma, flavor and nutritional characteristics during cheese making processes (Castro-Puyana et al., 2017). The most powerful sensory tool in cheese flavor research is descriptive sensory analysis. It requires trained human sensory evaluators to identify and quantify sensory aspects like appearance, aroma, flavor, texture… (Meilgaard, 1999). A good cheese flavor evaluator requires regular maintenance and 75-100 hours of training, which makes the descriptive analysis of flavor one of the most complex modalities to train (T. K. Singh et al., 2003). Furthermore, this form of sensory analysis can be challenged with subjectivity on the side of less trained or unprofessional panelists, the sensitivities of smell receptors (Bliss et al., 1996) and taste buds (Tuorila & Monteleone, 2009), since the sense of smell and taste varies with age, and in some cases, sex (Bliss et al., 1996; Peelle, 2019) and lifestyle activities such as smoking (Da Ré et al., 2018). Thus, human evaluation and inspection during food quality control may lead to inconsistent decisions. To cope with these challenges, metabolomics technology offers different tools that can identify and quantify the cheese’s flavors and aroma such as gas chromatography, mass spectrometry, aroma extract dilution analysis (AEDA) and odor activity value (OAV) (Pu et al., 2020). Gas chromatographic methods are widely applied in food science and technology due to an efficient compound separation and versatility. However, these metabolomics analyses demand expensive instrumentation and are time consuming as the optimum recovery of flavor compounds usually requires more than one procedure to avoid degradation and formation of artifacts and reach a detectable concentration of the components (Gomes et al., 2014; T. K. Singh et al., 2003; Weurman, 1969).

The final characteristics of a cheese, mainly flavors and aroma, are due to the complex dynamics and biochemical reactions involving the enzymes produced by cheese microorganisms and the milk components (lactose, fats, proteins). Therefore, analyzing these enzymatic profiles through sequencing methods such as metagenomics, proteomics and transcriptomics can serve to predict and characterize the metabolic profile of cheese. These sequencing methods are well developed and relatively cost-effective compared to the metabolomics experiments that directly measure the metabolic profiles of dairy products (Wang et al., 2023). Using an integrative approach to analyze 16S rRNA sequencing and metabolomics data collected from industrial and artisanal cheddar cheeses, Ashfari et al (Afshari et al., 2020) could detect strong relationships between the cheese microbiota and metabolome and uncovered specific taxa and metabolites that contributed to these relationships. Bertuzzi et al (Bertuzzi et al., 2018) have investigated the metabolic potential of the resident microorganisms of a surface-ripened cheese through whole-metagenome shotgun sequencing. They showed how variations in microbial populations influence important aspects of cheese ripening, especially flavor development. In human-based studies, Mallick et al (Mallick et al., 2019) have developed MelonnPan, an elastic net regularization method that can predict gut metabolites from the metagenomic data of gut microbial communities. This approach displayed promising performance and can be used for the prediction of metabolomes in similar studies where only microbiome is available. Similarly, neural networks have been developed to infer metabolic profiles from metagenomic and uncover microbe–metabolite relationships in environmental and clinical settings, such as soil biocrust wetting, cystic fibrosis and inflammatory bowel disease (Le et al., 2020; Morton et al., 2019; Reiman et al., 2021).

To our knowledge, the possibility of inferring cheese metabolic profile from microbial metatranscriptomics has not been sufficiently explored yet. In this work, we investigated the metatranscriptome-metabolome relationship in an experimental surface-ripened cheese by means of predictive models and correlation analyses. We trained Elastic Net (EN) and Random Forest (RF) models to infer the metabolic outcome of the cheese from its microbial gene expression profile. Despite the sparseness of the data, the accuracy of the models could reach 50 to 83 % in a completely independent validation set. Moreover, the analysis of the genes selected by the modeling procedure showed their consistency with biological pathways, mainly metabolic ones, that could be related to the observed outcomes. Our results demonstrate that metatranscriptomics data can be used as an informative proxy to estimate flavor profiles in cheese.

## Materials and Methods

### Extraction of RNA from cheese samples and rRNA depletion

RNA was extracted from rind sample aliquots without prior separation of microbial cells and total RNA was then subjected to rRNA depletion, as previously described in (Monnet et al., 2016)

### Library construction, RNA sequencing, mapping and processing

Library construction and RNA sequencing were performed by the Next Generation Sequencing Core Facility from the Institute for Integrative Biology of the Cell (Gif-sur-Yvette cedex 91198, France). Libraries were prepared with the Truseq SBS Kit v3 (50 cycles, FC-401-3002, Illumina), single read, 55 bp insert size. The sequencing reads were trimmed for the presence of Illumina adapter sequences using Cutadapt 1.3 (Martin, 2011), rRNA sequences were removed using SortMeRNA 4.2.0 (Kopylova et al., 2012), read sequences were mapped against the reference database using STAR 2.7.5a (Dobin et al., 2013). Raw data are available on ENA repository: https://www.ebi.ac.uk/ena/browser/view/PRJEB59186.

### Volatile compounds

Volatile compounds extraction and analyses were done as previously described in (Forquin et al., 2011). The volatile compounds were analyzed by a dynamic headspace analyzer (Purge and Trap HP 3547A; Agilent Technologies, Garches, France) fitted with a sorbent trap (Tenax) and a cryofocusing module. The concentrator was coupled to a gas chromatograph (GC G1530A; Agilent Technologies) connected to a mass spectrometer detector (MSD 5973; Agilent Technologies). Data were analysed with the software Agilent ChemStation and transformed from CDF raw files to mzml derived files with the software Lablicate OpenChrom. Aroma metabolites were identified by NIST 17 spectral library. Volatile quantification was performed using signal abundance (TIC - Total Ion Current). Raw data are available on the Metabolight repository: https://www.ebi.ac.uk/metabolights/MTBLS6968.

### Train and test transcriptomics and metabolomics datasets

Our data consist of longitudinal transcriptomics and metabolomics data collected from two independent experiments, that were used separately for training and testing our classification. Each of the training and test datasets contains one transcriptomics set (the predictors) set and one metabolomics set (the outcomes). The rows in the transcriptomics datasets represent time points and the columns represent sequenced genes. In the metabolomics data, rows represent the same time points and columns contain 6 measured metabolite classes: Alcohols, Aldehydes, Alkanes, Ketones, Esters and Sulphur compounds.

The training data were collected from an experiment with a surface-ripened cheese, fermented and ripened using a reduced microbial community composed of six bacterial and three yeast species: *Glutamicibacter arilaitensis* (Ga), *Brevibacterium aurantiacum* (Ba), *Corynebacterium casei* (Cc), *Hafnia alvei* (Ha), *Lactococcus lactis* (Ll), *Staphylococcus equorum* (Se), *Debaryomyces hansenii* (Dh), *Geotrichum candidum* (Gc) and *Kluyveromyces lactis* (Kl). There were three different experimental conditions: Control, and two other conditions where the Dh or the Gc yeast was omitted from the medium. There were three replicates at each of the <u>four</u> time points: Day 7, Day 14, Day 24 and Day 31.

The test data set is identical to the training set in terms of cheese type and microbial community, except for the experimental conditions (perturbation) which were set according to a variation in NaCl concentration (Dugat-Bony et al., 2015). There were three replicates at each of the <u>five</u> time points Day 1, Day 7, Day 14, Day 24 and Day 31.

### Normalization of the transcriptomics data

To reduce the biases induced by biological and technical artifacts, we adopted a method proposed by Klingenberg et al. (Klingenberg & Meinicke, 2017). The procedure relies on splitting the transcriptomics data by microbial species then, on each subset, performing the Trimmed Mean of the log expression ratios (M-values) normalization (TMM) (Robinson & Oshlack, 2010), which accounts for the sequencing depth and the RNA composition.

Both training and test transcriptomics data were split into nine species-specific datasets, each representing a given microbial species. In each species-specific dataset the lowly expressed genes, i.e, genes with less than 10 reads in three randomly selected samples, were filtered out. After this step, only two genes remained in *Staphylococcus equorum* data in the test set, thus, this species was discarded. As a result, we got nine and eight filtered species-specific datasets in the training and test set, respectively. Finally, the filtered and normalized species-specific datasets were merged to re-obtain the entire training and test datasets used for the modeling procedure.

### Orthology-based shrinkage of transcriptomics data

The metabolic profile in cheese is the result of the contribution of different enzymes expressed by the different microbes. Despite their differences, bacteria and yeasts can express proteins and enzymes having similar functions. These proteins (genes) are called orthologous proteins/genes (Gabaldón & Koonin, 2013). To reduce data dimensionality, we chose to sum together the expressions of the orthologous genes (same KEGG Orthology identifier (Kanehisa & Goto, 2000)). This allowed the shrinkage of the sample sizes in both training and test sets to almost the half (Table 1). This is a first dimensionality reduction step which we adopted before training predictive models. The second step is feature selection which is described below.

**Table 1:** Sample size (i.e. number of measured transcripts) of the training and test datasets before and after filtering out the lowly expressed genes and merging of the orthologous genes. The number of observations is 34 and 24 in the training and test set, respectively.

|  | Raw data | Filtered genes | Orthologous genes merged |
| --- | --- | --- | --- |
| Training set | 9868 | 8210 | 4245 |
| Test set | 3899 | 1769 | 1279 |

### Metabolomics data processing

Alongside gene expression, the amounts of six metabolite classes were measured in the two experiments: Alcohols, Aldehydes, Alkanes, Ketones, Esters, and Sulphur com- pounds. For each of the training and test sets, we had the corresponding metabolites amounts (observations) per each time point.

We trained classification instead of regression models, because the small number of available samples did not provide a sufficient representation of a range of continuous values. A bimodal distribution of values was considered reasonable after visual inspection (see Figure 1). The numerical values were converted into levels (categories) using k-means clustering (Macqueen, 1967) with two centers (*k* = 2).

**Figure 1:**
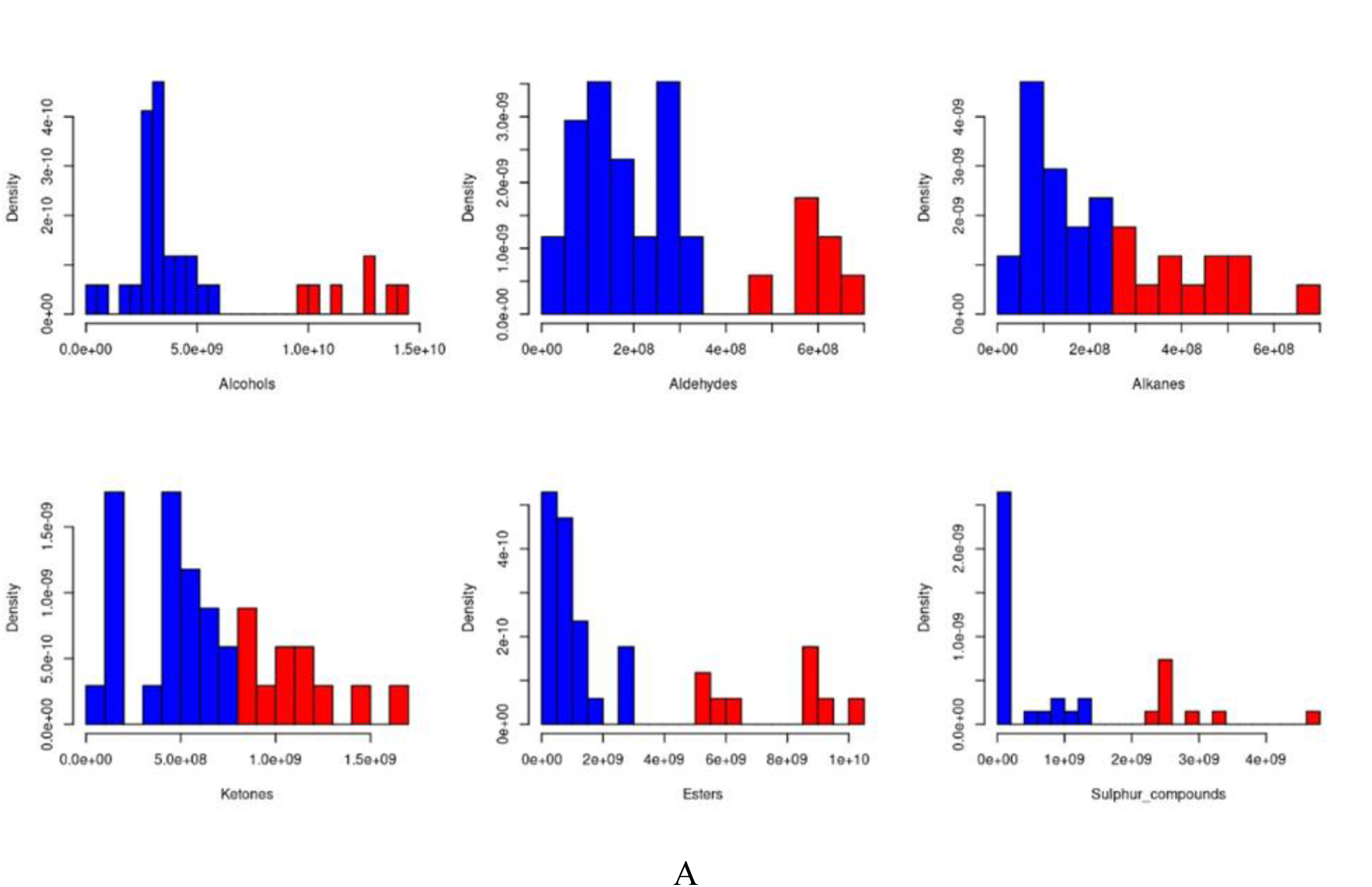
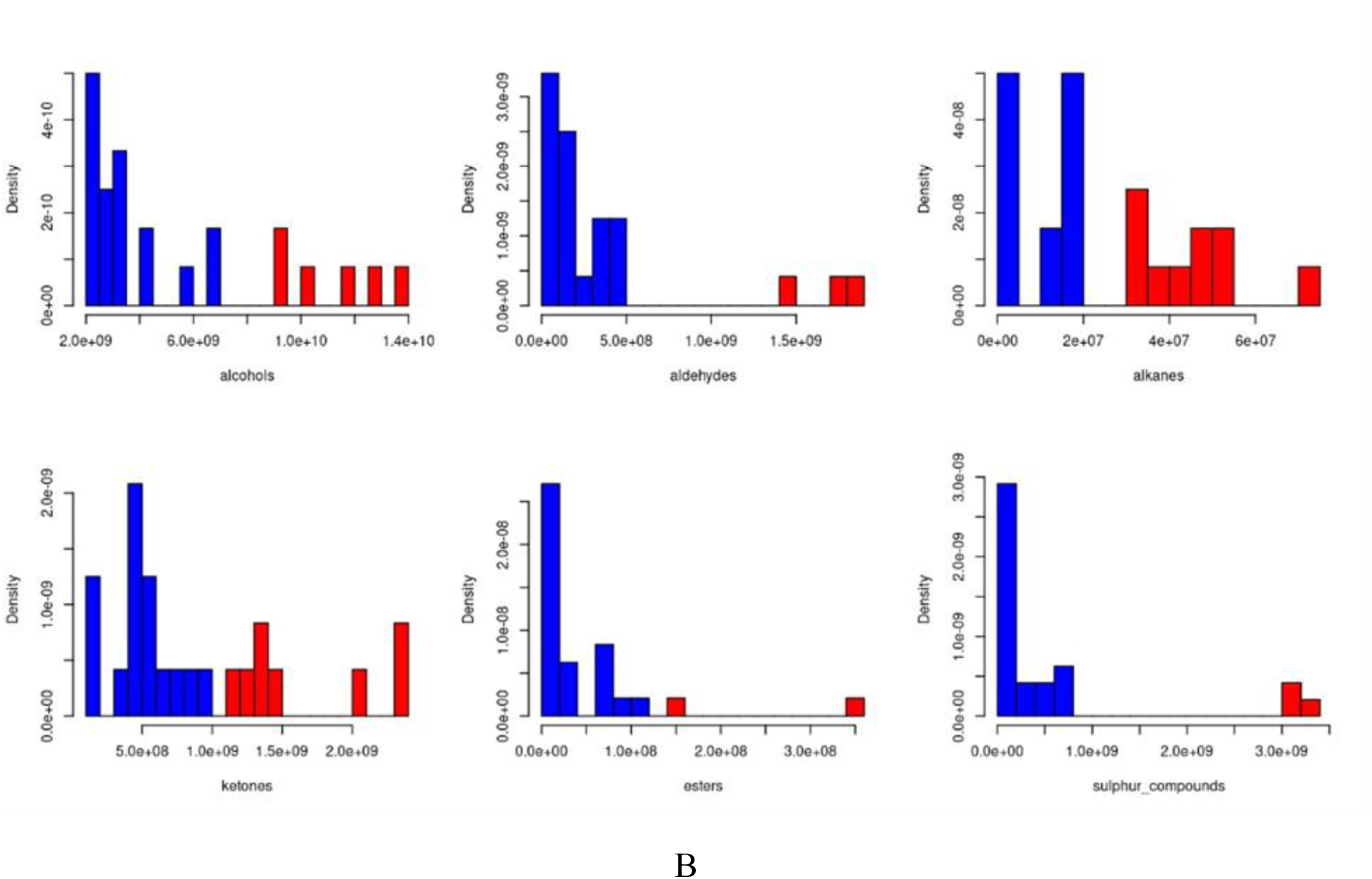
Distribution of the amount of each metabolite in the training (a) and test data (b). Blue and red represent the Low and High levels, respectively, determined by k-means clustering.

We labeled the continuous values in each cluster by “High” or “Low”. Each of the two data sets as well as the metabolite class were clustered and labeled separately. The original values of the metabolites amounts and the created labels are provided in the supplementary table 14.

### Training data and test data: further processing

Here, we describe the issues we had to deal with before being able to use these data for training and validation the predictive models.

#### 1. Same metabolites but different genes

The two transcriptomics datasets were collected from two similar but not totally identical experiments. The expression profiles of microbes depend on the conditions in which they grow, therefore, the number as well as the type of the expressed genes vary from a time point to another, and, from a condition to another.

Given that training and test transcriptomics data were collected using two different experimental designs, the genes found in these datasets don’t overlap totally. In the training set 4245 expressed genes were detected while in the test set there were 1279. The number of common genes between these datasets is 1062 genes (based on KEGG (Kanehisa & Goto, 2000) Orthology Numbers). The problem that arises here is that we can’t train a model on a given set of features then validate it in a different set, even if the predicted variable (metabolite class) is exactly the same. So we had two options: 1) training the models and cross-validate them using only the training dataset (FTS, 4245 genes and 34 observations); 2) Training the models on a reduced training set (RTS) which is a subset of FTS containing only the 1062 common genes with the test set, and then validate the models on the independent test set (Table 3). Option 1 is good in terms of completeness of predictor variables, since all the microbial transcriptomic profile is explored by the modeling procedure to find the best fitting, but on the other hand the validation is less robust since it is a cross-validation on the same data set. Option 2 offers a more robust validation, as the test set is independent from the training set, but almost half of the predictive features (genes) are not explored, and this may result in a less accurate estimation of the transcriptome-metabolome correlation in cheese.

**Table 3:** The size of the original training (FTS) and test sets, in addition to the created Reduced Training Set.

| | Training Set (FTS) | Test Set | Training Set ∩ Test Set | Reduced Training Set |
| --- | --- | --- | --- | --- |
| Genes | 4245 | 1279 | 1062 | 1062 |
| Samples | 34 | 24 | - | 34 |

To guarantee both completeness and robustness, we performed two modeling procedures in parallel: 1) We trained and cross-validated the classification models using FTS consisting of 4245 genes/34 samples and, 2) We trained the models using RTS (1062 common genes/34 samples) then validated them on the independent test set.

#### 2. The data are high-dimensional cursed and imbalanced

Let’s remind that both FTS and RTS are dimensionally high, i.e, there are more predictors (genes) than observations (levels of metabolites). There are 34 data points (metabolite levels) in FTS and RTS, and 4145 and 1062 genes, respectively. Secondly, we investigated the levels of the six metabolite classes and we found a class imbalance, i.e, the “High” class has less samples than the “Low” one (Table 4). When the response variable (classes) is skewed, it can break down relatively robust procedures used for unskewed data (Chen & He, 2013). To handle the high dimensionality and class imbalance, we performed EN-based feature selection followed by RF classifiers to cope with both problems.

**Table 4:**
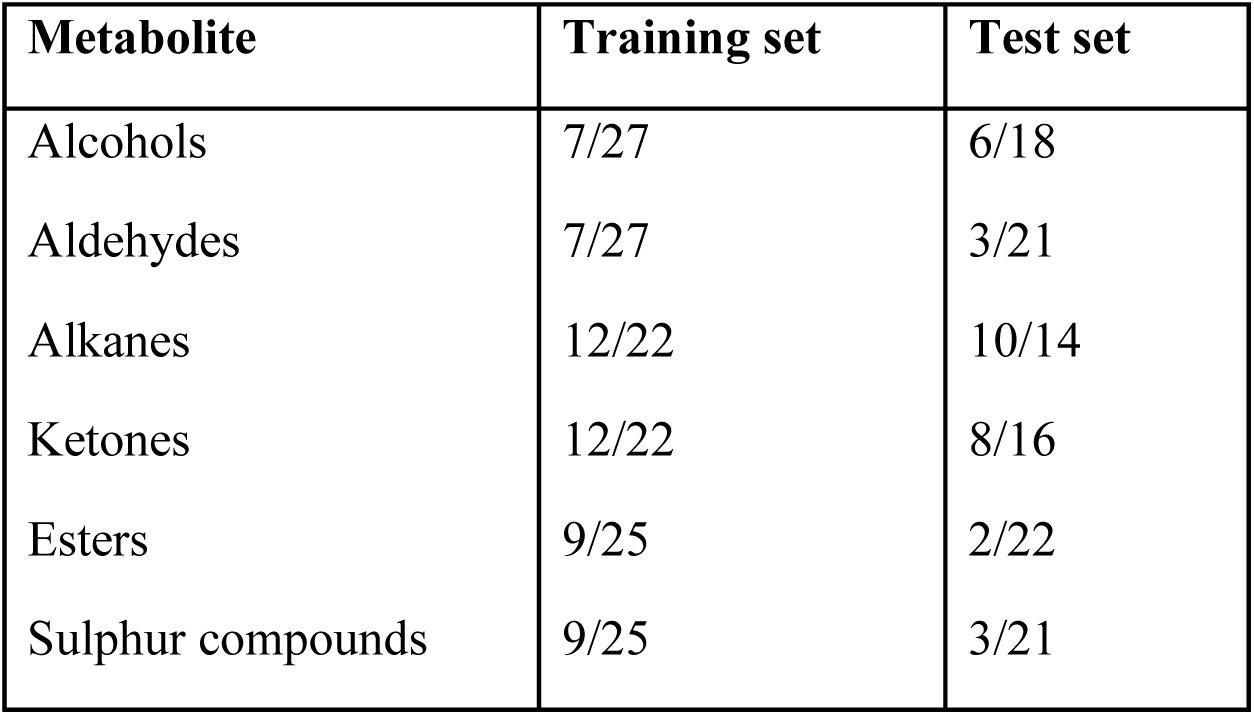
The number of High/Low labels associated to each numerical value of the metabolite amounts.

### Elastic Net as a feature selection algorithm

As stated earlier, we wanted to train classification models that predict the Low/High levels of a given metabolite class, based on microbial gene expression values. More precisely, we trained six different models, each one predicting one metabolite class. In a similar work, Mallick et al. (Mallick et al., 2019) have trained elastic net regularization model to predict gut metabolites from the metagenomics data of human gut microbial communities. In our work, we adopted a similar methodology. EN combines Ridge and LASSO features, i.e, it handles multicollinearity (predictors correlation) and it performs feature selection by assigning null coefficient to non-informative predictors (Biernat & Lutz, 2015; James et al., 2013). To run the EN algorithm, we used the *caret (Kuhn, 2008)* and *glmnet (Hastie, 2021)*R packages.

### RF models trained on EN-selected features

To deal with both high dimensionality and class imbalance, we trained RF classifiers on the features (genes) selected from FTS and RTS. A RF model can yield noticeably different performance on the same dataset when the random seed changes as this alters the bootstrap samples and feature selections that define the ensemble. Therefore, to effectively assess our modeling procedure, we run 1000 replications with 1000 different random seeds. As a result, we got 1000 similar but not identical RF models then we averaged their performances to get an overall estimation (Table 8).

### Biological validation of the selected features

To assess the biological relevance of the genes (predictors) selected by EN, we performed a Spearman correlation test (Spearman, 1961) between their eigengenes and the amounts of each metabolite class. The eigengene E is the first principal component summarizing the expression profiles in each selected gene set (Langfelder & Horvath, 2008). Then, we performed a permutation test to see how significant is the EN-feature selection as compared to a random gene selection.

Secondly, the selected genes have been analyzed to assess their relevance and association to biological pathways through over-representation analysis (ORA), using the *enrichKEGG()* function from *clusterProfiler* R package (Yu et al., 2012). As metabolic pathways are interconnected and often overlap (Judge & Dodd, 2020), i.e, enzymes and metabolites of a specific reaction chain can be also involved in another reaction chain, we merged selected predictors of the six models to construct the gene set used to run the ORA.

### Assessment of the classification models

For a more accurate estimation of the models’ performances, we used metrics suitable for imbalanced learning: Balanced Accuracy, Area Under ROC curve (AUC), F1 score and Specificity. Accuracy is the simplest metric used for the evaluation of predictive models. It is the proportion of good predictions out of all the predictions. Accuracy is widely used, however it can be misleading in the case of imbalanced classes. Therefore, we used *Balanced Accuracy* as an alternative as it takes into account both the proportions of correctly predicted positive (Sensitivity or recall) and negative classes, then averages them. Area Under the Receiver Operating Characteristic Curve (AUC) indicates how well the model discriminates between “Low” and “High” classes. *AUC=0.5* indicates that the models’ prediction are random. *AUC >* 0.7 indicates that the model has fair to excellent (AUC = 1) discrimination ability (Çorbacoğlu & Aksel, 2023). F1-measure represent the harmonic mean of *Precision* and *Recall*.

The RF models trained on the FTS were assessed using a Leave-One-Out cross-validation (LOO-CV), whereas those trained on the RTS were assessed using the independent test set. The performances of the 1000 replicates were averaged to get an overall estimation. RF model fitting were performed using the R package *caret* v 6.0.94 (Kuhn, 2008).

### Analysis of microbes’ contribution to the selected features

The selected gene sets from both FTS and RTS were analyzed to define which gene(s) is expressed by which microbe(s). Each organism’s contribution is represented by a percentage:

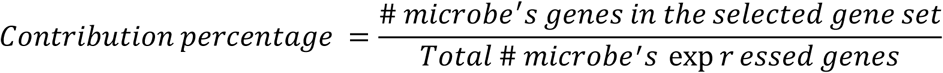

Microbes highly involved in metabolic process leading to flavor formation should have high contribution percentages, i.e, the feature selection step should capture more genes expressed by those microbes. To check this assumption, we measured the relationship between the different microbial species and the metabolites classes. We computed a Spearman correlation (Spearman, 1961) between the growth rate of these microbes (colony forming units) and the measured amounts of metabolites across the samples.

## Results

### Performances of the RF models

All the models trained and crossvalidated on FTS performed very well in terms of accuracy (BA [0.82, 0.94]) as well as in prediction of the less abundant class “High” (Specificity [0.67, 0.89]) (Table 9). These are promising results; however, the LOO cross-validation is not so robust as the models were trained and tested using the same dataset (FTS).

**Table 9:**
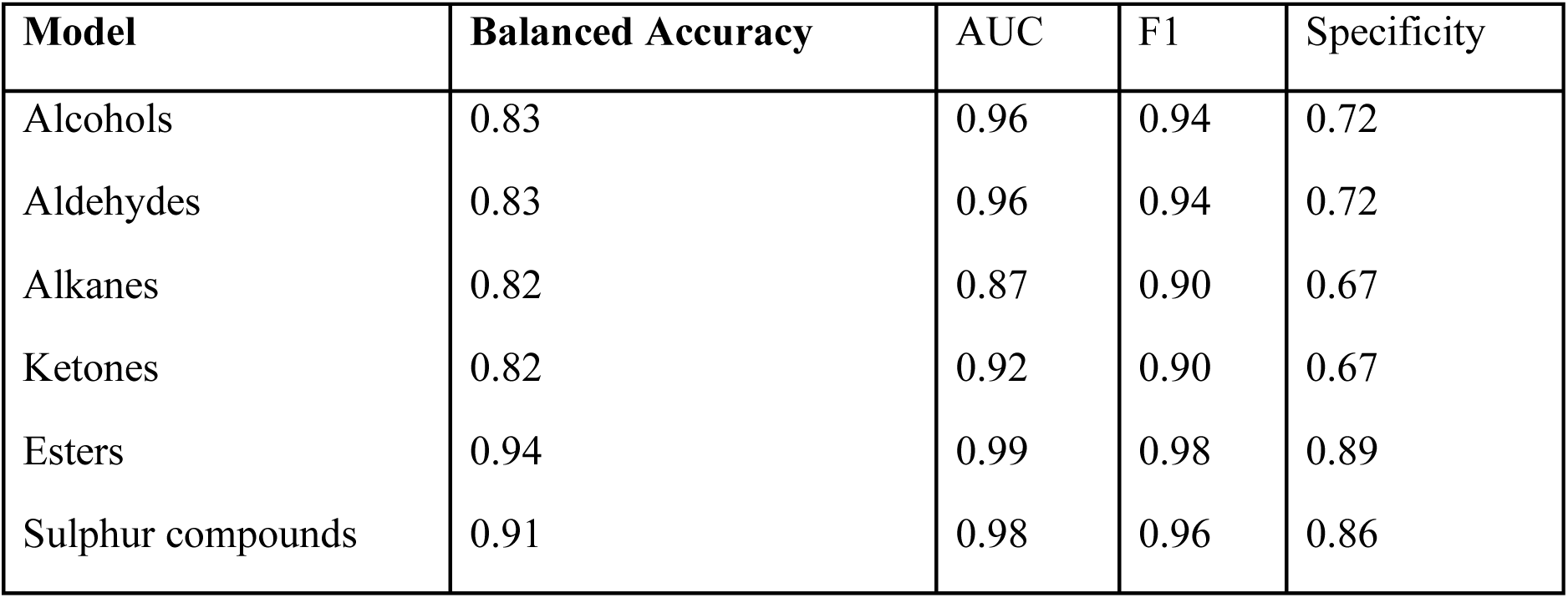
Performances of the RF classifiers trained after EN-feature selection on FTS and cross-validated using LOO.

| Model | Balanced Accuracy | AUC | F1 | Specificity |
| --- | --- | --- | --- | --- |
| Alcohols | 0.83 | 0.96 | 0.94 | 0.72 |
| Aldehydes | 0.83 | 0.96 | 0.94 | 0.72 |
| Alkanes | 0.82 | 0.87 | 0.90 | 0.67 |
| Ketones | 0.82 | 0.92 | 0.90 | 0.67 |
| Esters | 0.94 | 0.99 | 0.98 | 0.89 |
| Sulphur compounds | 0.91 | 0.98 | 0.96 | 0.86 |

In the second case in which the RF were trained using the predictors selected from the RTS then validated on the independent test set, all models showed rather good accuracy (BA [0.49, 0.83]). Alcohols, Aldehydes and Esters models had good specificity scores, 0.76, 0.77 and 1, respectively, while the specificity score for the remaining models ranged from 0 to 0.40. The Sulphur compounds model failed to predict any “High” class (less abundant class) (Table 10).

**Table 10:** Performances of the RF classifiers trained after EN-feature selection on RTS and validated using the independent test set.

| Model | Balanced Accuracy | AUC | F1 | Specificity |
| --- | --- | --- | --- | --- |
| Alcohols | 0.83 | 0.92 | 0.91 | 0.76 |
| Aldehydes | 0.79 | 0.84 | 0.87 | 0.77 |
| Alkanes | 0.49 | 0.54 | 0.58 | 0.40 |
| Ketones | 0.49 | 0.52 | 0.79 | 0.007 |
| Esters | 0.58 | 0.92 | 0.27 | 1 |
| Sulphur compounds | 0.50 | 0.44 | 0.93 | 0 |

### Correlation between the selected predictors and the biological traits

The EN features selection has retained about 0.004-0.03 % and 0.007-0.08 % of the total number of genes in FTS and RTS, respectively. To assess the correlation between the selected predictors and the metabolites classes, each predictors set has been represented by its eigengene E (the first component vector), then Spearman correlation has been computed between E and the corresponding metabolite class (Table 11). In both FTS and RTS, E correlates well with the corresponding metabolite class where the highest correlation was observed for Sulphur compounds and the lowest for Alcohols.

**Table 11:** Number of features selected in each modeling procedure and the correlation of their eigengene with the metabolites classes. *: the correlation is not better than random (pvalue > 0.05).

| Model | # Selected genes from FTS | Correlation with the metabolite | # Selected genes from RTS | Correlation with the metabolite |
| --- | --- | --- | --- | --- |
| Alcohols | 45 | 0.57 | 23 | 0.51 |
| Aldehydes | 45 | 0.66* | 23 | 0.68* |
| Alkanes | 17 | 0.79 | 7 | 0.76 |
| Ketones | 48 | 0.79 | 28 | 0.74 |
| Esters | 128 | 0.85* | 82 | 0.77* |
| Sulphur compounds | 42 | 0.92 | 25 | 0.92 |

Despite the strong correlations between aldehydes and esters and the corresponding selected predictors, they are not significantly higher than correlations obtained by a random predictor selection. In the case of Aldehydes, 20 % and 15 % of the random correlations were equal or higher than 0.66 and 0.68, respectively. For Esters, 23 % and 42 % of the permuted correlations were equal or higher than 0.85 and 0.77, respectively. This could be due to the high dimensionality of the data which causes the model to be unstable, i.e, it can find several patterns that explain well the metabolite-genes relationship.

Secondly, to assess the biological relevance of the selected predictors from both FTS and RTS, we run an over-representation analysis to see which biological pathways were significantly represented. Six metabolic pathways were associated to the predictors selected by the EN algorithm (Table 12): Microbial metabolism in diverse environments, Phenylalanine metabolism, Carbon metabolism and Citrate cycle (TCA cycle). Propanoate metabolism was exclusively enriched in FTS whereas Pyruvate metabolism was exclusively enriched in RTS (Table 12). Cheese ripening, mainly, the development of aromatic compounds (alcohols, aldehydes, ketones…) involve many metabolic pathways such as carbon metabolism and citrate cycle (Fox et al., 2004; McSweeney, 2004).

**Table 12:** KEGG pathways over-represented using the genes selected from FTS and RTS.

| Over-represented Pathways | FTS | RTS |
| --- | --- | --- |
| Microbial metabolism in diverse environments | ✓ | ✓ |
| Phenylalanine metabolism | ✓ | ✓ |
| Carbon metabolism | ✓ | ✓ |
| Citrate cycle (TCA cycle) | ✓ | ✓ |
| Propanoate metabolism | ✓ | ✓ |
| Pyruvate metabolism |  |  |

Esters play an important role in the flavor profiles of various cheeses, contributing with fruity and sweet characteristics (McSweeney & Sousa, 2000). Several selected genes are associated with pyruvate, glycolysis, and lactate metabolism. Acetate esters, which are commonly found in cheese, are formed via a reaction between acyl-CoA and an alcohol (Richoux et al., 2008). Genes linked to pyruvate metabolism within the ester signatures (e.g., K00163 and K00382) are involved in the production of acetyl-CoA, that can be utilized for ester formation. Amino acid biosynthesis pathways also influence ester formation, as indicated by the production of phosphoserine esters (K00831) within the serine biosynthesis pathway. Notably, serine can be converted to cysteine, a precursor to major volatile compounds in cheese.

Aldehydes in cheese originate from pyruvate metabolism, lipid oxidation, and amino acid catabolism, with the latter being the most relevant for flavor development. Amino acid catabolism is particularly important in mold- and smear-ripened cheeses, where the breakdown of free amino acids produces intermediates like amines and α-keto acids that are subsequently converted into aldehydes. These aldehydes are further metabolized into alcohols or acids, contributing to the complex flavor profile of cheese (Eugster et al., 2000). EN selected aldehyde dehydrogenase (K04021) and alcohol dehydrogenase (K00121) which are involved in the conversion of aldehydes to acetyl-CoA and alcohols respectively. Furthermore, proteins such as K03695 involved in amino acid degradation, highlight the role of free amino acids in aldehyde production. The model also identified multiple proteins from the *eut* operon which convert ethanolamine into acetaldehyde, a yogurt, green and nutty aroma in cheese (Smit et al., 2005).

Genes selected in Ketone model were mainly from lipid catabolism, glycerol catabolism, amino acid biosynthesis and aromatic compound metabolism. They impart a blue-cheese-like flavor in cheese. Ketone signatures support the role of lipolysis in ketone formation. Initially, lipase enzymes hydrolyze triglycerides into free fatty acids (FFAs), di- and mono-glycerides, and glycerol, which would explain the presence of a glycerol dehydrogenase (K00005) and acyl CoA oxidase (K00232) in the signatures. Acyl CoA oxidase could also be involved in methyl ketone formation (Walker & Mills, 2014). FFAs can be oxidized to β-ketoacids, and then decarboxylated to corresponding methyl ketones (McSweeney & Sousa, 2000). Other genes were related to methionine and threonine metabolism (K00789, K00872, K01243). Ketones are tightly related to amino acids due to their conversion to alpha keto acids through transamination. Moreover, Brevibacteria, present in the current study, contains methionine-γ-lyase, that catalyzes a reaction to make methanethiol and α-ketobutyrate from methionine. Threonine is also deaminated to produce α-ketobutyrate and propionic acid (Ganesan & Weimer, 2017). These compounds are important intermediates in the production of aromatic compounds in cheese.

In smear and ripened cheese, sulfur compounds contribute greatly to the characteristic aroma of cheese, giving it a sulfurous, cheesy and garlicky flavor. They are synthesized from the degradation of the sulfur containing amino acids methionine and cysteine, and they inlcude: hydrogen sulfide, methanethiol, dimethyl sulfide dimethyl disulfide trisulfide, S-methyl thioesters (Landaud et al., 2008; Sourabié et al., 2012). K00789, an S-adenosyl-L-methionine (SAM) synthase, is part of the methionine degradation pathway. It is a methyl donor and is therefore central in many biological processes. In the context of cheese, SAM can be converted into S-methylmethionine, a precursor to dimethyl sulfide. SAM also participated in pathways that produce homocysteine, a precursor to hydrogen sulfide. Additionally, the K01243 gene, encoding adenosylhomocysteine nucleosidase, is involved in methionine salvage pathway. The selection of this feature by the model demonstrates the importance of methionine in aromatic compound production. Taken together, the genes selected by the models give an indication of the contribution of each metabolite class to the organoleptic characteristics of cheese. In Table 13 we report some of the genes encoding enzymes involved in metabolic pathways.

**Table 13:**
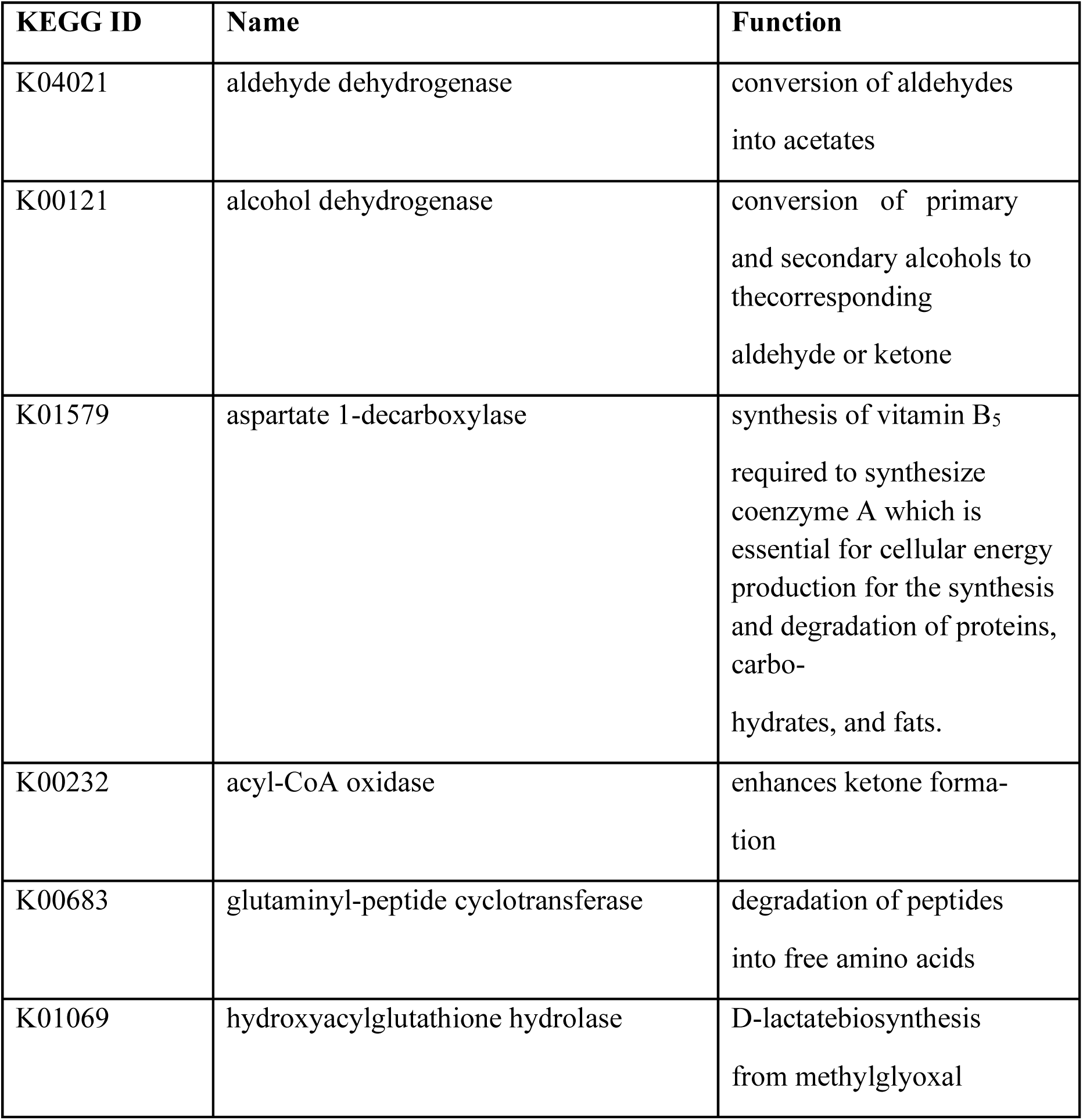
Non-exhaustive list of the genes selected by the EN algorithm and their biochemical functions.

### Microbes’ contribution to the signatures

The comparison of the bacterial and yeast expression profiles in both training and test transcriptomics data showed that yeasts express about two to three times more genes (mRNAs) than bacteria: it was thus expected that the yeasts would contribute more to the models’ signatures. However, the EN feature selection has retained more bacterial than yeast genes as shown in figure 2. In FTS-selected gene set, the percentages of selected genes from each bacteria were: *Ga* 13 %, *Ba* 20 %, *Cc* 10 %, *Ha* 10%, *Ll* 11 %, *Se* 13 %, *Dh* 5 %, *Gc* 5 % and *Kl* 6 %. This holds true also in RTS where bacteria contributed more than yeasts to the selected gene set: *Ga* 30 %, *Ba* 41%, *Cc* 27 %, *Ha* 26 %, *Ll* 24 %, *Dh* 14 %, *Gc* 11 % and *Kl* 16 %. In both selection procedures, Ba species had the highest contribution. Note that the percentages don’t’ sum up to 100 because each microbe’s percentage is independent from the others, i.e. the number of selected genes belonging to each microbe was divided by the total number of its expressed genes, and not by the total number of selected genes (see Methods).

**Figure 2:**
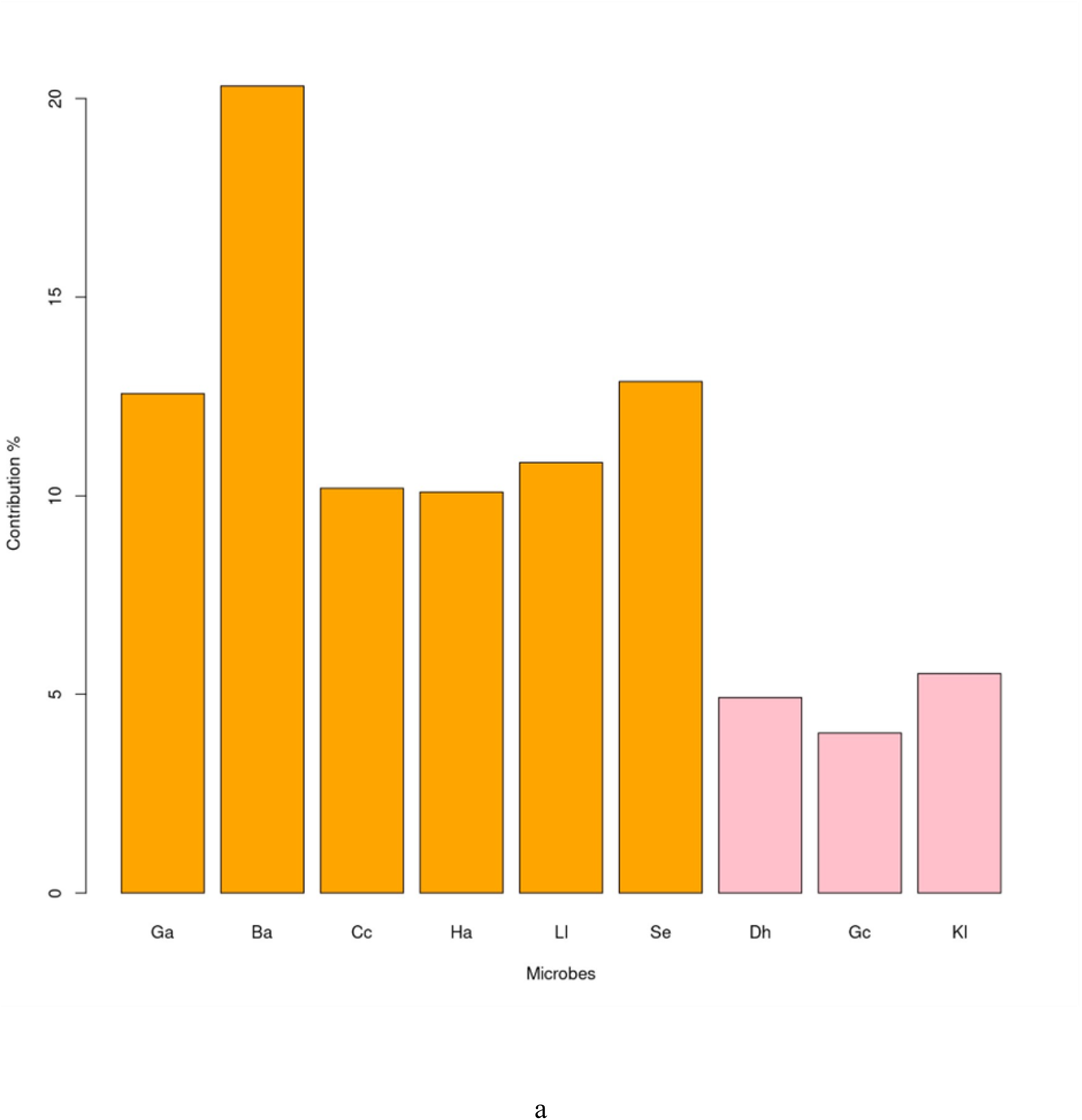
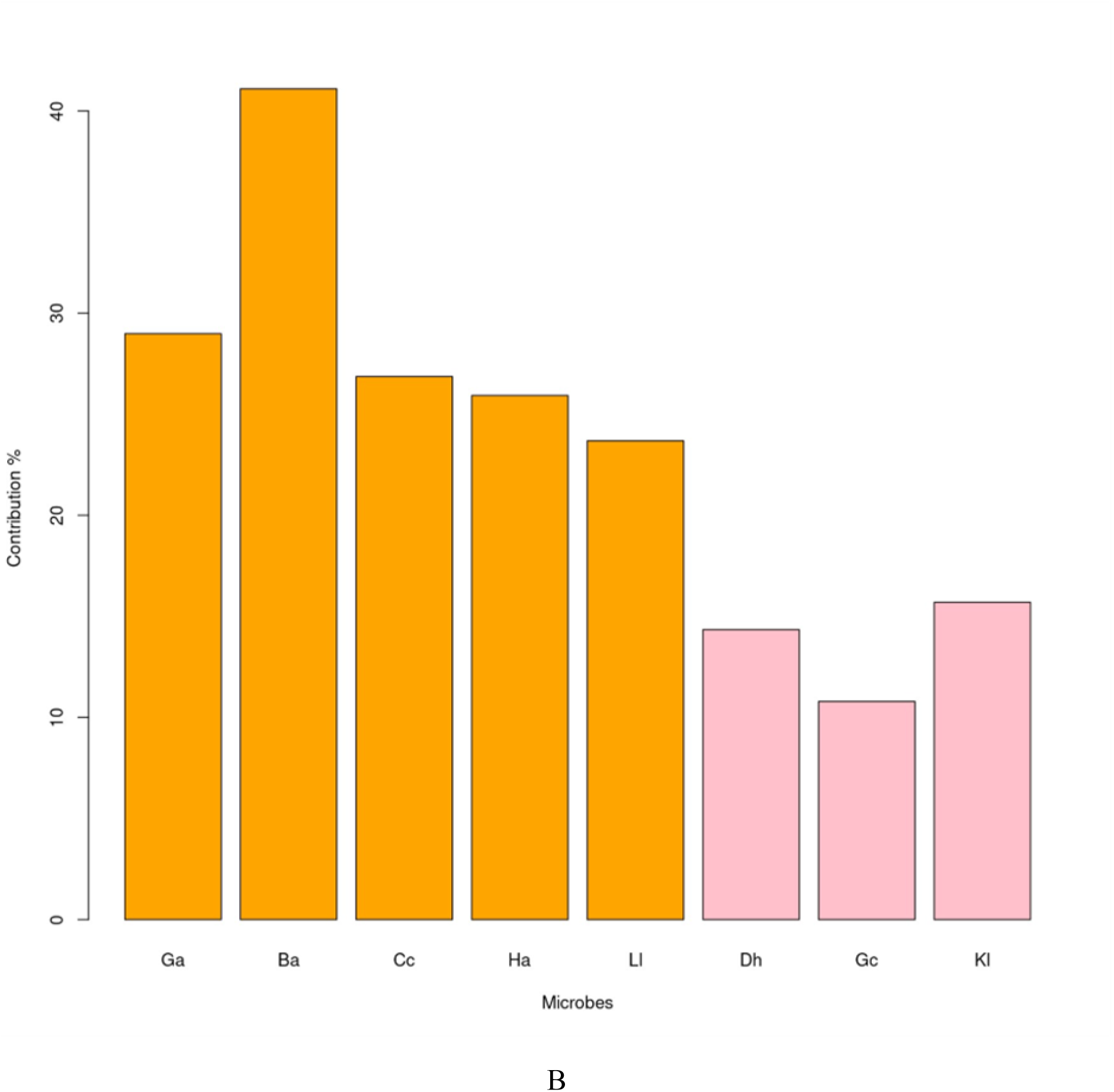
Contribution percentage of each species to the selected gene sets from both FTS (a) and RTS (b). Orange: bacteria. Pink: yeasts.

This suggests that, in this specific case, i.e, experimental cheese, bacteria contribute to flavor formation more than yeasts. These observations are in line with the correlation results (Figure 3) where flavor amounts were more correlated with the growth of bacteria than yeasts. In other words, the increase or decrease of bacterial metabolic activity due to a decreases/increase of their number, induces a significant variation of the cheese flavor profile.

**Figure 3:**
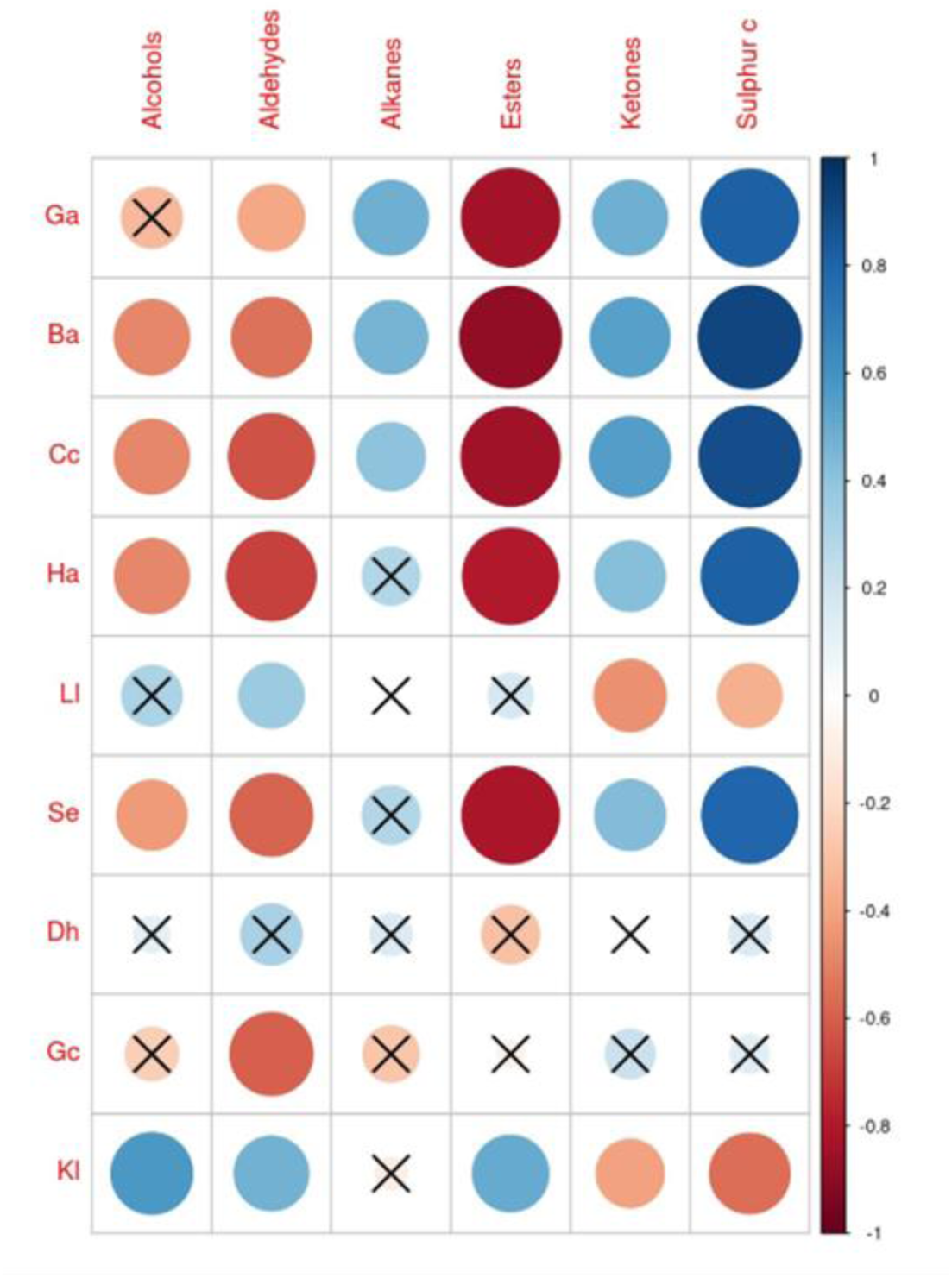
Spearman correlations between microbial growth and metabolites amount. Black crosses represent non-significant correlations. The areas of the circles and the colors’ intensity are proportional to the absolute value of the correlation coefficients.

## Discussion

One of the objectives of this study was to demonstrate that estimation of cheese flavor profiles from the gene expression of the microbial communities involved in cheese processing is feasible. Ultimately, cheese quality control (monitoring metabolic/flavor profile) could be complemented by such straightforward *in silico* predictions, which are more cost-effective and less time consuming than the traditional sensory and metabolomics analyses techniques. This can reduce cheese quality control and cheese making costs, especially for large-scale cheese/food manufacturers that are very aware of customer preferences in terms of quality and affordability.

Despite the small dataset size and the imbalance between high and low flavour labeling classes, the performances of the cross-validated models were high, with an accuracy ranging from 82% to 94%. Furthermore, we also implemented and analysis with a more robust validation using a totally independent test set, and also in this case the performances were overall satisfying (AUC *>* 0.5, except for Sulphur compounds model). It’s true that most of them suffered from a low specificity, as it often happens with imbalanced classes (Ferri et al., 2009; Japkowicz, 2013), but on average their accuracy can be qualified as good.

## Acknowledgements

D. R., D. S., A. M. and R. M. acknowledge EU MSCA-ITN-ETN “E-MUSE” Project n. 956126

